# Forest Dormouse (*Dryomys nitedula*) populations in southern Italy belong to a deeply divergent evolutionary lineage

**DOI:** 10.1101/210070

**Authors:** R. Bisconti, G. Aloise, A. Siclari, V. Fava, M Provenzano, P. Arduino, A. Chiocchio, G. Nascetti, D. Canestrelli

## Abstract

The Forest Dormouse (*Dryomys nitedula*) is a small rodent with a wide, albeit severely fragmented distribution, ranging from central Europe to central Asia. Within the Italian region, *D. nitedula* populations are restricted to forested mountain areas of two largely disconnected regions, the eastern Alps and the Calabria region, where two distinct subspecies (*D. nitedula intermedius* and *D. nitedula aspromontis*, respectively) have been described on the basis of phenotypic characters (i.e., fur colour). Here we analysed *D. nitedula* samples from both regions, to investigate patterns of genetic divergence and phylogenetic relationship among these two populations. Genetic variation was studied at the level of one mitochondrial (cytochrome b gene) and three nuclear gene fragments (exon1 of the interstitial retinoid-binding protein, exon 10 of the growth hormone receptor, and recombination activating gene 1). Phylogenetic analyses were performed using Maximum Likelihood and Bayesian inference methods. *D. n. aspromontis* and *D. n. intermedius* were found to be reciprocally monophyletic in all the phylogenetic analyses, and the genetic divergence observed between them at the mitochondrial *CYTB* gene was conspicuous (HKY: 0.044) when compared to previously observed values among many sister species of rodents. Our results clearly show that *D. nitedula aspromontis* is a deeply divergent, narrow endemic evolutionary lineage, and its conservation needs should be carefully evaluated in the near future. Moreover, such deep genetic divergence, together with phenotypic differentiation between *D. n. intermedius* and *D. n. aspromontis,* suggest that *D. nitedula* populations in southern Italy might belong to a distinct, previously unrecognized species.

## Introduction

The Italian Peninsula has long been identified as a major component of the Western Mediterranean biodiversity hotspot, and as an important glacial refugium for temperate animal species throughout the Plio-Pleistocene (Hewitt, 2011). The advent and extensive application of genetic markers to the study of geographic variation have much improved our understanding of key biogeographic patterns and historical processes within this area, revealing expansion-contraction dynamics, population fragmentations into multiple Pleistocene refugia, hidden hybrid zones, as well as the occurrence of a plethora of cryptic and deeply divergent evolutionary lineages (Barbanera et al., 2009; Canestrelli et al., 2006a, 2006b, 2007a, 2007b, 2008a, 2010, 2012a, 2012b, 2014a, 2014b; Canestrelli and Nascetti, 2008; Castiglia et al., 2007, 2016; Colangelo et al., 2012; Grill et al., 2009; Kindler et al., 2013; Lecocq et al., 2013; Lo Brutto et al., 2010; Louy et al., 2013; Maura et al., 2014; Mezzasalma et al., 2015; Nascetti et al., 2005; Salvi et al., 2013; Salvi et al., 2017; Simonsen and Huemer, 2014; Wauters et al., 2017).

The Forest Dormouse *Dryomys nitedula* (Pallas, 1778) is a small rodent with a wide, albeit fragmented geographic distribution, ranging from eastern and southern Europe to central Asia (Krystufek and Vohralik, 1994). Despite its wide distribution, current knowledge about its ecology and systematics is still scanty. The species has arboreal and nocturnal habits, and it has been observed from the sea level to above 2000 m a.s.l., within a wide variety of habitats, but with marked differences found among local populations (Krystufek and Vohralik, 1994; Paolucci et al., 1989; Amori et al., 2008). Together with the wide but geographically structured variation in body size, coat colour, and to a lesser extent morphology, these differences among local populations have led several authors to suggest possible occurrences of cryptic species within *D. nitedula* (Holden, 2005). Although a comprehensive investigation of its molecular systematic is still missing, the few data available seem to support this hypothesis (e.g. Grigoryeva et al., 2015), and indicate that cryptic divergent lineages may exist within this nominal species.

Within the Italian Peninsula, *D. nitedula* populations are restricted to forested mountain areas of two largely disconnected regions: eastern Alps and southern Italy (Aspromonte, Sila, and Pollino mountain massifs). However, this large distributional gap could have been narrower in the recent past. Fossil data suggested that the species occurred in central Italy, at least before the last glacial phase (65-35 thousand years ago; see Kotsakis, 1991, 2003). Based on differences in coat colour patterns (Nehring, 1902; Von Lehmann, 1964), the two populations from the eastern Alps and southern Italy have so far been described as two distinct subspecies *D. nitedula intermedius* Nehring, 1902 and *D. nitedula aspromontis* Von Lehmann, 1964 respectively, with the latter showing a brighter grey fur and a distinctive white spot on the tip of the tail (Von Lehmann, 1964). In spite of extensive faunistic surveys in the Calabria region (Aloise and Cagnin, unpublished data), *D. n. aspromontis* individuals have hither to been found only at altitudes above 1000 m a.s.l., and only within beech (*Fagus sylvatica*) dominated forests (Cagnin and Aloise, 1995), whereas along the Alps, the species has also been observed at lower altitudes, and mostly within mixed forests of broadleaf trees and conifers (Paolucci et al., 1989). However, cytogenetic and morphometric differences have not been observed between both subspecies (Civitelli et al., 1995; Filippucci et al., 1995), and a limited genetic differentiation have been reported based on preliminary allozyme data (Filippucci et al., 1995), leading to uncertainty about to the correct taxonomic assignment of the populations in southern Italy (Amori et al., 2008).

In this study, we investigate patterns of genetic divergence between *D. n. aspromontis* and *D. n. intermedius* by analysing patterns of sequence variation at the level of one mitochondrial and three nuclear gene fragments. Our aim was to better characterize the phylogenetic relationships between the forest dormouse population in southern Italy and its conspecific populations in the north. In fact, given the large geographic gap among the subspecies, dispersal and gene exchange look rather implausible. Consequently, assessing whether *D. n. aspromontis* can be better defined as a marginally differentiated geographical isolate or as a unique evolutionary lineage might have major implications, not only for taxonomy but also of profound conservation value.

## Materials and Methods

In total, 15 samples of *D. nitedula* were analysed (see Figure 1 and Table 1). Tail-tip samples of *D. n. aspromontis* (n = 8) were collected in the field since this species uses tail autotomy as an anti-predator behaviour (Mohr, 1941). Alternatively, tissue samples were also picked up from road-killed individuals. All samples were transported to the laboratory and stored in 95% ethanol until DNA extraction. Tissue samples of *D. n. intermedius* (n = 7) were kindly provided by the Science Museum of Trento (Muse) and Padova University as ethanol preserved specimens (see Table 1).

**Table 1.** Geographic location of the 15 samples of *Dryomysnitedula*analysed in this study.

| Sample | Source | Code | Locality | Longitude (E) | Latitude (N) | Altitude (m) |
| --- | --- | --- | --- | --- | --- | --- |
| 1 | This study | ASP1 | Montalto | 15.908 | 38.159 | 1825 |
| 2 | This study | ASP2 | Tre Limiti | 15.859 | 38.137 | 1600 |
| 3 | This study | SL1 | Macchiad'Orso | 16.646 | 39.129 | 1624 |
| 4 | This study | SL2 | Monte Gariglione | 16.642 | 39.132 | 1669 |
| 5 | This study | SL3 | Monte Gariglione | 16.642 | 39.132 | 1669 |
| 6 | This study | SL4 | Monte Gariglione | 16.642 | 39.132 | 1669 |
| 7 | This study | SL5 | Monte Gariglione | 16.642 | 39.132 | 1669 |
| 8 | This study | SL6 | Monte Gariglione | 16.642 | 39.132 | 1669 |
| 9 | Science Museum of Trento | MTSN 1120 | ForestaDemaniale di Cadino | 11.403 | 46.190 | 1650 |
| 10 | Science Museum of Trento | MTSN 1121 | ForestaDemaniale di Cadino | 11.406 | 46.191 | 1600 |
| 11 | Science Museum of Trento | MTSN 1122 | ForestaDemaniale di Cadino | 11.406 | 46.191 | 1600 |
| 12 | Science Museum of Trento | MTSN 1123 | ForestaDemaniale di Cadino | 11.389 | 46.214 | 1850 |
| 13 | Science Museum of Trento | MTSN 1124 | ForestaDemaniale di Cadino | 11.390 | 46.217 | 1800 |
| 14 | University of Padova | ALB 2379 | Val di Fiemme | 11.738 | 46.306 | 1510 |
| 15 | University of Padova | ASD4 | Asiago | 11.516 | 45.967 | 1700 |

**Figure 1.**
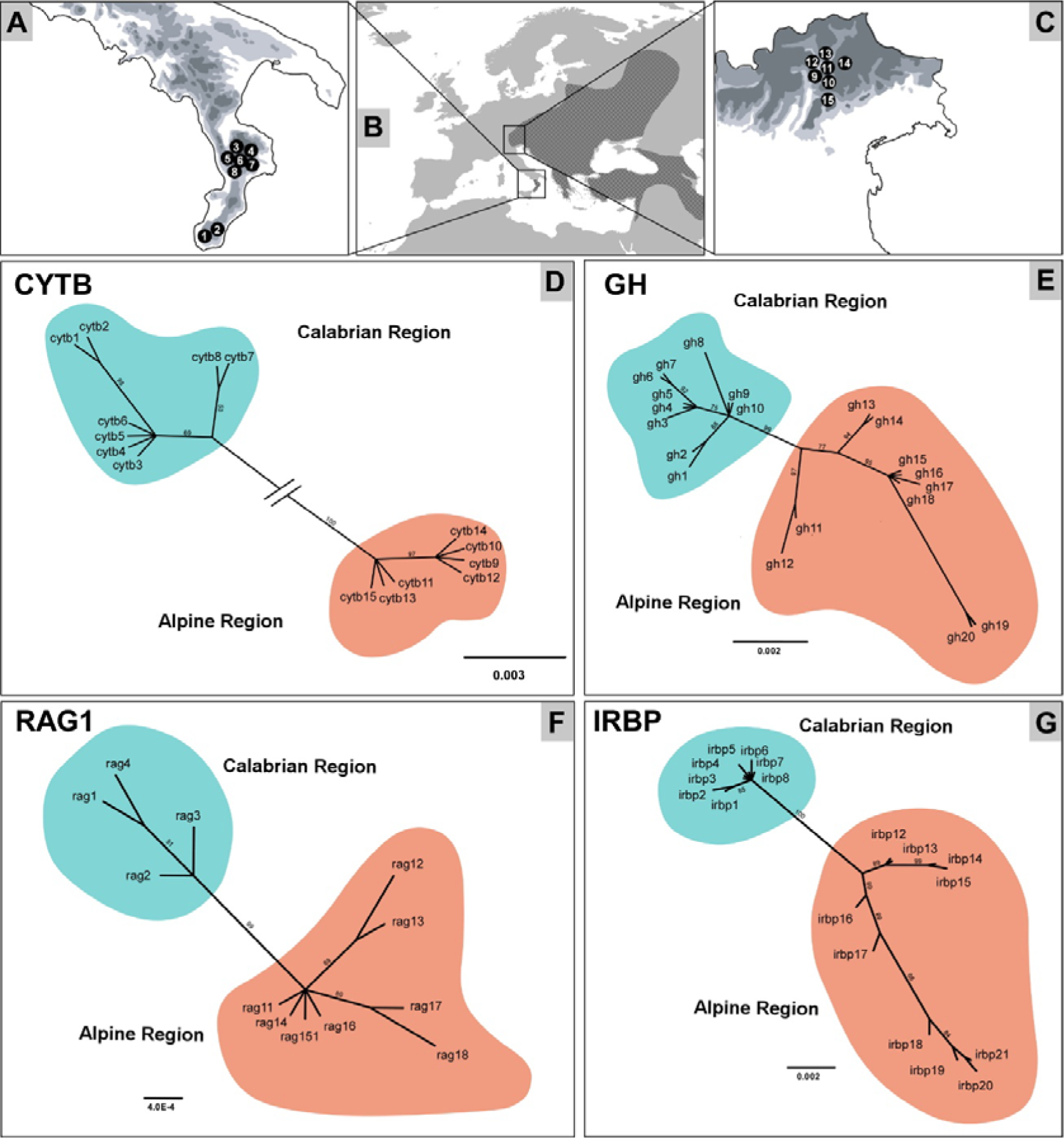
**A)** Geographic location of the *Dryomys nitedula aspromontis* samples analysed for the present study; localities are numbered as in Table 1. **B)** Geographic distribution of *Dryomys nitedula* in Europe and neighbouring regions (redrawn from Juškaitis, 2014). **C)** Geographic location of the *Dryomys nitedula intermedius* samples analysed for the present study; localities are numbered as in Table 1. **D-G)** Phylogenetic trees inferred for the four gene fragments analysed based on a Bayesian inference procedure; numbers indicate Bayesian posterior probabilities.

**Figure 2.**
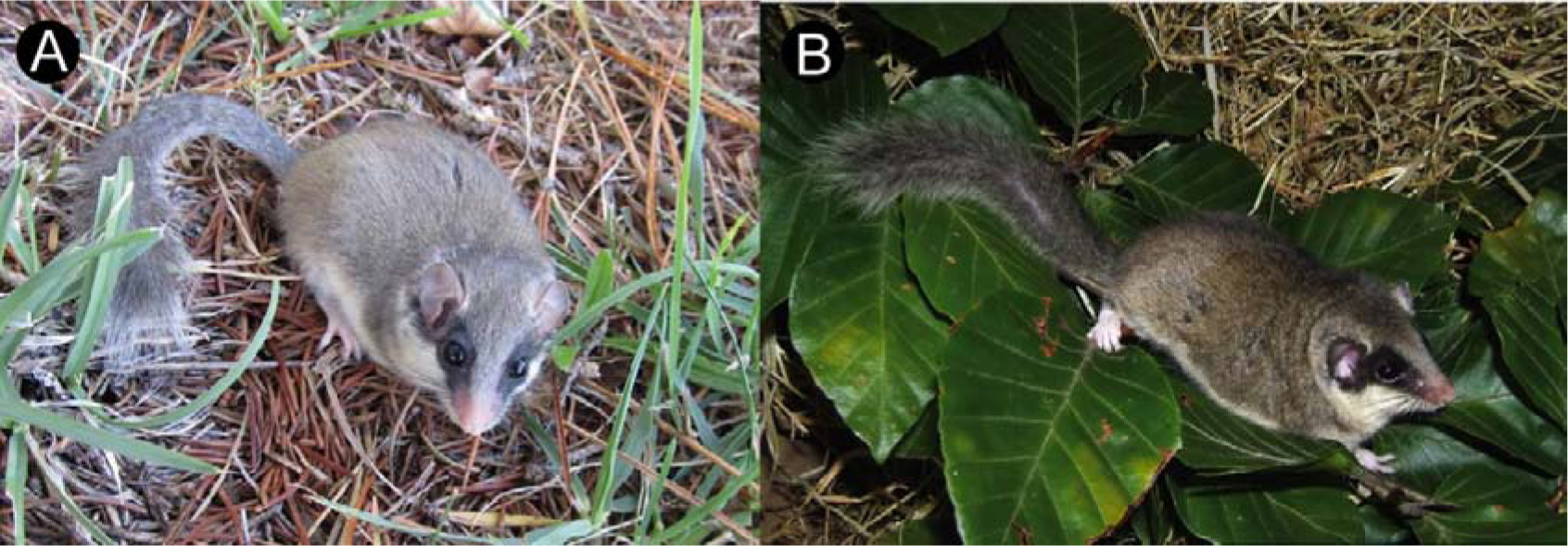
Pictures of the two subspecies of *Dryomy snitedula* inhabiting the Italian peninsula. **A)** *Dryomys nitedula aspromontis* (Monte Altare Longobucco; Photo credit: A. Pellegrino). **B)** *Dryomy snitedula intermedius* (Photo credit: L. Lapini).

Whole genomic DNA was extracted using ZR universal kit (Zymo Research), following the standard DNA extraction protocol provided. Partial mitochondrial sequences of the cytochrome b gene (*CYTB*) were obtained using the following primers (Grigoryeva et al., 2015): F_Dr.n_cyt (TGACAAACATCCGTAAAACT) and R_Dr.n_int (AAAAGCGGGTTAGTGTTGC). Amplifications by polymerase chain reaction (PCR)were performed with modifications from the original protocol (Grigoryeva et al., 2015): an initial denaturation step at 94°C for 3 minutes, followed by 30 repeated cycles of 94°C for 30 seconds, 54°C for 30 seconds and 72°C for 1 minute, and a single final step at 72°C for 5 minutes. Three nuclear gene fragments were amplified: exon1 interstitial retinoid-binding protein (*IRBP*), exon 10 of the growth hormone receptor (*GHR*), and a portion of recombination activating gene 1 (*RAG1*). PCR primers used and cycling conditions were the same as presented in Pisano et al., 2015. Amplifications were carried out using identical PCR mixtures for all gene fragments analysed, including: 20 ng of extracted DNA in a 25-µL reaction mix containing MgCL2 (2.5 mmol/L), the reaction buffer (1X; Promega), four dNTPs (0.2 mmol/L each), two primers (0.2 µmol/L each), and the enzyme Taq polymerase (1 unit; Promega). PCR products were purified and sequenced by Macrogen Inc. (htpp://macrogen.com) using the ABI PRISM 3700 sequencing system.

The sequences obtained were visually checked by using CHROMAS 2.31 (TechnelysiumLtd.), and they were aligned with CLUSTALX (Thompson et al., 1997) with the default settings. All the sequences obtained were deposited in the GenBank database (accession numbers: XXX-XXX [to be populated upon acceptance]). Sequences diversity and divergence patterns among sequences were evaluated using DIVEIN (Deng et al., 2010). Nuclear heterozygous sequences were phased using PHASE 2.1 (Stephens et al., 2003) with the default options, whereas the occurrence of recombination was assessed using the pairwise homoplasy index (PHI statistic, Bruen et al., 2006) in SPLITSTREE v.4.11 (Huson and Bryant, 2006).

The best-fit model of sequence evolution was selected for each analysed gene fragment among 88 alternative models using the Bayesian Information Criterion (BIC) in JMODELTEST 2.1.3 (Darriba et al., 2012). This method suggested HKY as the best substitution model for the mitochondrial fragment (*CYTB*), HKY+I for the *IRBP* gene and JC+I for the *GHR* and *RAG1* genes.

Phylogenetic trees were estimated by means of the Maximum-Likelihood (ML) algorithm as implemented inPhyML program (Guidon et al., 2010), using default settings for all parameters, with the following exceptions: i) node support was assessed through a non-parametric bootstrap procedure based on 1000pseudo-replicates ii) the best substitution model, as indicated by JMODELTEST, was used for each analysed marker. To check for consistency among different phylogenetic tree estimation procedures, phylogenetic trees were also estimated based on the Bayesian inference procedure (BI) by the MRBAYES v.3.2.1 software (Ronquist et al., 2012). For this purpose, four Monte Carlo Markov chains were run for 10 million generations with trees sampled every1000 generations, and the first 25% of the resulting trees discarded as a burn-in.

## Results

For all the individuals analysed we obtained sequences of length 427 bp for the *CYTB* gene fragment, 889 bp for *GHR*, 1216 for *IRBP*, and 826 bp for *RAG1.*The 427 bp mitochondrial region *CYTB* showed 21 variable positions, 20 parsimony informative, whereas no indels, stop codons, and nonsense codons were observed. The *GHR* gene showed 18 variable positions of which 17 parsimony informative, the *IRBP* gene presented 29 variable positions of which 24 parsimony informative, and the *RAG1* gene showed 8 variable positions of which 6 parsimony informative. The PHI test carried out with the nuclear gene fragments did not suggest statistically significant indications of recombination events.

Since phylogenetic trees inferred by means of ML and BI methods yielded fully congruent tree topologies, only results based on BI will be presented here (ML trees available upon request). As shown in Figure 1, for all the genetic markers analysed, tree topologies clearly identified samples belonging to *D. n. aspromontis* (southern Italy) and *D. n. intermedius* (north-eastern Italy) as two reciprocally monophyletic and well supported lineages, with no instances of common haplotype. Mean sequence divergence between haplotypes within each group was minimal, and below values observed between groups at all the markers analysed (see Table 2). The highest value of divergence estimated between both groups (HKY = 0.044; p-distance = 0.043) was observed at mtDNA gene fragment (*CYTB*).

**Table 2.** Mean sequence divergence (maximum-likelihood estimate) within and between the main groups of haplotypes recovered by the phylogenetic analyses carried out among the *D. nitedula* samples analysed in the present study. Standard errors are given in brackets.

| Level of variation | Geographic region | *CYTB* | *GHR* | *IRBP* | *RAG1* |
| --- | --- | --- | --- | --- | --- |
| *Within groups* | North-eastern Italy | 0.001 | 0.006 | 0.007 | 0.002 |
|  |  | (0.000) | (0.001) | (0.001) | (0.000) |
|  | Southern Italy | 0.003 | 0.002 | 0.001 | 0.002 |
|  |  | (0.000) | (0.000) | (0.000) | (0.000) |
| *Between groups* |  | 0.044 | 0.009 | 0.013 | 0.005 |
|  |  | (0.000) | (0.000) | (0.000) | (0.000) |

## Discussion

Studies of intraspecific diversity within the forest dormouse have almost entirely been based on phenotypic patterns of variation (but see e.g. Filippucci et al., 1995; Grigoryeva et al., 2015), and lead to the description of several subspecies within the nominal species *D. nitedula*. However, to what extent these phenotypic variants are in fact evolutionary independent lineages still remains largely unknown. In this study, we analysed the patterns of genetic divergence between the geographically isolated populations of forest dormouse in southern Italy (*D. n. aspromontis*), and their geographically closest population in north-eastern Italy (*D. n. intermedius*).

Our results clearly show that *D. n. aspromontis* is an independent evolutionary unit, monophyletic at all the markers analysed, and deeply divergent at the mtDNA from geographically the closest population in north-eastern Italy (HKY=0.044). These results seem to contradict with data from Filippucci et al. (1995), in which a rather low level of allozymic differentiation (D=0.03) was suggested. Nevertheless, while some discordance in terms of genetic diversity and differentiation patterns would not be surprising (Toews and Brelsford, 2012), a direct comparison between the two divergence estimates would be hardly meaningful. Indeed, given the fully allopatric distribution of the two lineages, a discussion of the possible discordance could only be based on a comparison of genetic distance metrics derived from distinct methodological approaches. Nonetheless, it is worth noting that while Filippucci and colleagues (1995) did not identify a single allozymic locus of fully diagnostic value between the subspecies, our results indicated a perfect reciprocal monophyly at the three nuclear loci studied, thus suggesting a lack of power resolution of the allozymic loci used by Filippucci and colleagues (1995).

During the data analysis, we refrained from using mtDNA for a molecular dating exercise because incomplete taxon sampling might strongly affect the resulting estimates (Poux et al., 2008; Nabhan and Sarkar, 2012), and our samples of *D. n. intermedius* was largely incomplete. Nevertheless, we cannot fail to notice that the sequence divergence observed at the *CYTB* between the northern and southern samples suggested a much older divergence for *D. n. aspromontis* than the mid-Holocene (approximately 10,000 years) or Late Pleistocene (35,000-65,000) as previously hypothesized based on morphological and fossil data, respectively (Roesler and Witte, 1968; Krystufek and Vohralik, 1994; Filippucci et al., 1995). In fact, using the mutation rate of 0.0217 mutations/site/million years recently estimated for the *CYTB* in mammals (Igea et al., 2015), the amount of sequence divergence we found suggested a divergence time between the subspecies around 1 million years ago (i.e. the Early Pleistocene), thus predated this event compared to previous estimates. Consequently, the single fossil record of *D. nitedula* found in central Italy (Kotsakis, 1991, 2003), might suggest a recent range contraction into southern Italy of a formerly ‘peninsular’ lineage, as already shown for a large amount of animal species in the area, (e.g. Canestrelli et al, 2006a; 2008; Grill et al., 2009; Castiglia et al., 2016; Colangelo et al., 2012) rather than a very recent (i.e. Late Pleistocene to mid-Holocene) colonization of southern Italy from the Alps as previously thought (Roesler and Witte, 1968; Krystufek and Vohralik, 1994; Filippucci et al., 1995).

Our results have major implications for forest dormouse conservation in southern Italy. In fact, our results definitely identify this lineage as a unique evolutionarily significant unit (ESU, sensu Moritz, 1994), endemic to this geographic area and, to the state of knowledge, fragmented into three geographic isolates restricted to mountain tops above 1000 m a.s.l. in the Aspromonte, Sila, and Pollino mountain massifs. Further research is needed to assess the demographic consistency and patterns of genetic diversity of these isolates, and to better define the most appropriate management strategy of this narrow endemic lineage.

Finally, our results could also have a major taxonomic implication that might be critical for conservation and management, since priorities in conservation strategies are defined based on species status and species diversity (see e.g., the Convention on International Trade in Endangered Species of Wild Fauna and Flora (CITES) listed species; the IUCN red list of threatened species). Assigning allopatric populations to the species or subspecies rank based on the amount of genetic divergence might be problematic because patterns of reproductive isolation cannot be assessed in the field. Nevertheless, given the major theoretical and applied implications linked to the taxonomic rank, several attempts have been made in this regard either by exploring alternative definitions of the species conceptor scanning literature for plausible thresholds values of genetic divergence to assign a taxon to the species rank (for a perspective on mammals, see Baker and Bradley, 2006). In the case of *D. n. aspromontis*, the *CYTB* sequence divergence we found with respect to the closest population in north-eastern Italy, equals or even exceeds those observed among many sister species of mammals, and rodents in particular (see e.g. Michaux et al., 2002; Baker and Bradley, 2006; Wauters et al., 2017). Furthermore, *D. n. aspromontis* shows distinct morphological features, concerning unique coat colour pattern (see above). Accordingly, populations of the forest dormouse in southern Italy could in fact be assigned the species rank. In this case, *Dryomys aspromontis* Von Lehmann, 1964 would be available as the taxon name with a suitable common name as the Calabrian forest dormouse, since to the state of knowledge its current range is mostly restricted to this region. However, a note of caution is needed in the present case based on at least one major argument. The patterns of genetic diversity have not been investigated yet in *Dryomys nitedula* at the level of its entire range. Since there are several morphologically defined units (i.e. subspecies) stemming in geographical contiguity to one another within *Dryomys nitedula* from continental Europe to central Asia, a thorough examination of the associated patterns of genetic divergence and, most importantly, reproductive isolation might provide comparative yet important knowledge, in order to make better informed decisions about the correct taxonomic ranking of the southern Italian lineage as well.

## Acknowledgment

We thanks to Luca Lapini and A. (Sasà) Pellegrino for kindly sharing photos, as well as Maria Chiara Deflorian (Science Museum of Trento, Muse), Paolo Paolucci (Padova University), and Antonio Mazzei (University of Calabria) for their help with sample collection and/or with laboratory procedures. This research was supported by grants from the Italian Ministry of Education, University and Research (PRIN project 2012FRHYRA), and from the Aspromonte National Park.

